# The mammalian forelimb diversity as a morphological gradient of increasing evolutionary versatility

**DOI:** 10.1101/2022.07.26.501575

**Authors:** Priscila S. Rothier, Anne-Claire Fabre, Julien Clavel, Roger Benson, Anthony Herrel

## Abstract

Vertebrate limb morphology often reflects the environment, due to variation in locomotor requirements and other ecological traits. However, proximal and distal limb segments may evolve differently to each other, reflecting an anatomical gradient of functional specialization that has been suggested to be impacted by the timing of bone condensation during ontogeny. Here we explore whether the temporal sequence of bone condensation predicts variation in the capacity of evolution to generate morphological diversity between proximal and distal forelimb segments across more than 600 species of mammals. Our findings are consistent with the hypothesis that late developing distal limb elements should display greater morphological variation than more proximal limb elements, which condense earlier during morphogenesis. Distal limb elements, belonging to the autopod, not only exhibit higher diversity of form, but are also more integrated and, on average, show greater evolutionary versatility than intermediate and upper limb segments. Our findings indicate that the macroevolutionary patterns of proximal and distal limb segments are not the same, suggesting that strong functional selection, combined with the higher potential of development to generate variation in more distal limb structures, facilitate the evolution of high autopodial disparity in mammals.

## Introduction

The evolutionary origin of limbs marks the onset of the adaptive radiation of Tetrapoda (Shubin et al., 1997). From delicate wings to powerful excavating claws, from slender hooved legs to wide flattened flippers, limb formation is intrinsically integrated with and constrained by the determination of the tetrapod body plan (Raff, 1996). The tetrapod limb is typically composed of three basic components: the proximal stylopod (upper arm and thigh), the intermediate zeugopod (lower arm and calf), and the distal autopod (hand and foot). The proximal to distal organization of segments is correlated with their respective evolutionary appearance, the stylopod being the first structure to evolve, later followed by the zeugopod, and finally the autopod (Shubin et al., 1997). Although the three-segment pattern is conserved among quadruped tetrapods, the morphology of these structures along the proximo-distal axis may evolve differently among groups (Cooper et al., 2011; Galis et al., 2001; Holder, 1983; Sears et al., 2007).

Mammalian limbs are often studied for their exceptional morphological and ecological diversity, particularly in the forelimbs (see Figure 1; e.g., Polly 2007; Chen and Wilson 2015; Weaver and Grossnickle 2020; Howenstine et al. 2021; Lungmus and Angielczyk 2021). The forelimb is present in all mammal species and is typically more variable than the hind limb, possibly due to its greater number of functional roles (e.g., Polly, 2007; Schmidt and Fischer, 2009). In mammalian adult morphologies, the meristic composition of forelimb segments varies along the proximo-distal limb axis, where the autopod exhibits most of the diversity in terms of the number and position of skeletal elements (i.e., fusion and loss of carpal and tarsal bones and alteration of the phalangeal formula; Cooper et al., 2007; Hamrick, 2001; Holder, 1983; Luo et al., 2015; Saxena et al., 2017). In contrast, structures from the proximal segments are always present in the mammalian forelimb, displaying some but less frequent cases of element reduction and partial fusion of the zeugopod bones (observed in bats, manatees, horses, etc., Holder, 1983; Sears et al., 2017, 2007). Although this meristic information is useful to quantify major evolutionary changes in element composition, most of the morphological variation observed in mammalian limbs results from changes in the shapes and relative sizes of individual elements (i.e. variation of form) without changing the numbers of elements, and is often associated with functional adaptation (Fabre et al., 2013, 2015; Janis & Martín-Serra, 2020; Lungmus &,Angielczyk, 2021; Sears et al., 2017). Despite its importance, it remains unclear how the variation of form is partitioned between more proximal and distal skeletal elements.

**Figure 1.**
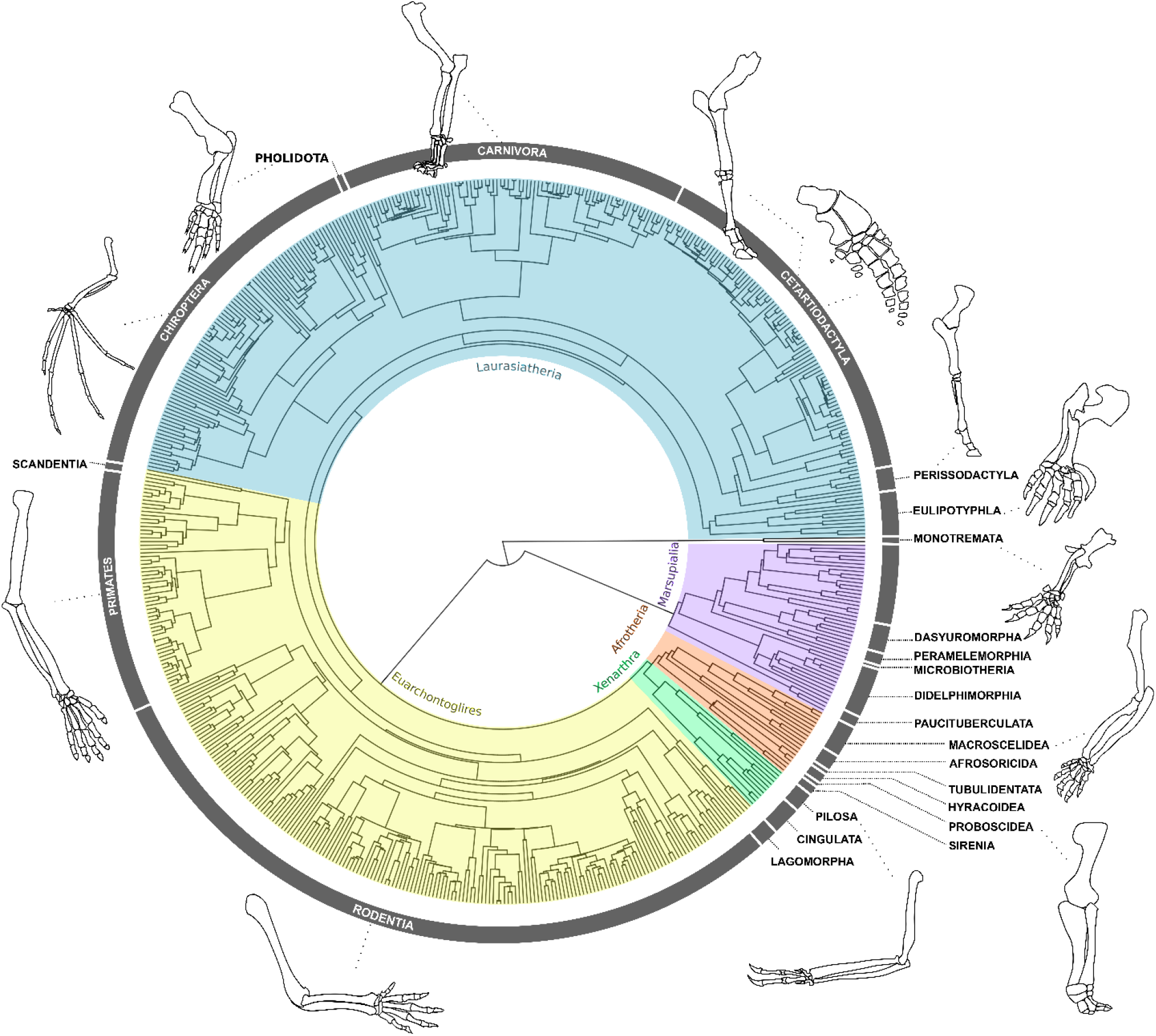
Forelimb diversity of mammals. The topology includes all genera examined in this work, representing the outstanding forelimb morphological variation for some of the species analysed. The topology was estimated using maximum clade credibility from a posterior sample of 10,000 trees published by Upham et al. (2019).

Both functional and developmental factors may predict that distal elements should show greater variation of form than more proximal elements. Developmental mechanisms predict this pattern due to the timing and spatial pattern of morphogenesis. Each limb initiates as a bud that extends from the body wall, where skeletal elements are generally specified in a proximal to distal sequence that matches to their evolutionary appearance during tetrapod origins: development begins with the stylopod, followed by the zeugopod, and terminating in the autopod at the distal end (Schneider and Shubin, 2013; Shubin et al., 1997; Stopper and Wagner, 2005).

Studies of mammals have revealed that different species have more similar forelimb morphology during early development, and become more disparate during later stages of morphogenesis (Ross et al., 2013). Likewise, gene expression is more conserved during early phases of limb development, compared to later phases (Maier et al., 2017). This has been shown for species with dramatically different adult limb morphologies: bats, pigs, opossums and mice (Maier et al., 2017), and these patterns might reflect the intrinsic temporal properties of embryogenesis (Galis et al., 2001; Maier et al., 2017; Sears et al., 2017). Specifically, early developmental processes mediating the initial specification of structures are generally more constrained than those governing later events, such as organ specialization (Kalinka and Tomancak, 2012). Therefore, because limb development proceeds proximo-to-distally, developmental perturbations at later phases may tend to accumulate higher morphological variation in distal elements (Hallgrímsson et al., 2002).

One way to infer the levels of developmental and functional constraints on adult morphologies is by quantifying the phenotypic integration among traits, inferred from the covariation between structures. Because the fore and hind limbs are serially homologous elements, they share genetic and developmental processes that give rise to strong phenotypic integration between and within the limbs (Ruvinsky and Gibson-Brown, 2000; Young and Hallgrímsson, 2005). The correlation between homologous limb segments of the fore- and hindlimbs (i.e., humerus with femur, radius with tibia, metacarpal with metatarsal) has been described for some mammalian groups, suggesting that proximal segments are highly integrated to each other (Hallgrímsson et al., 2002; Schmidt and Fischer, 2009; Young and Hallgrímsson, 2005). In contrast, more distal elements, of the hand and foot show more variable patterns of integration, which may reflect the accumulation of variation during later phases of development (Hallgrímsson et al., 2002; Rolian, 2009; Young and Hallgrímsson, 2005). Furthermore, because the autopod is the structure that interacts directly with the substrate, this limb segment is likely to experience more dynamic selective pressures favouring locomotor specialization in certain environments compared to proximal segments. A consequence for limb evolution is that the patterns and pace of morphological evolution might not be the same between proximal and distal segments.

Here, we investigate the evolutionary patterns underlying the morphological diversification of mammalian forelimb segments along a proximal-to-distal axis, using a comprehensive data set of 638 species, capturing over 85% of Mammalia family-level diversity (Table S1 of SI 1). We ask to what extent is the temporal structure of proximo-distal bone condensation consistent with the macroevolution of limb segment morphologies. We examined the diversification of limb skeletal elements by quantifying morphological diversity and integration using linear measurements of four forelimb bones (Figure 2, Table S2 of SI 2). We also estimated the macroevolutionary patterns of these elements using multivariate phylogenetic comparative models. First, we quantified the morphological diversity of each segment, testing the hypothesis that distal bones are morphologically more diverse than the proximal structures. Next, we investigated whether the strength of within-segment integration differs between proximo and distal limb elements. We predicted that proximal elements would be more integrated than distal ones, due to their earlier condensation during development. Finally, we inferred the macroevolutionary patterns for bones belonging to all limb segments, predicting positive associations between the temporal sequence of bone condensation and the capacity for evolution to generate morphological diversity. To our knowledge, this is the first time that the evolutionary patterns observed in the form of proximal versus distal limb elements are investigated using a broad phylogenetic and ecological sample of mammalian diversity, essential to address these questions.

**Figure 2.**
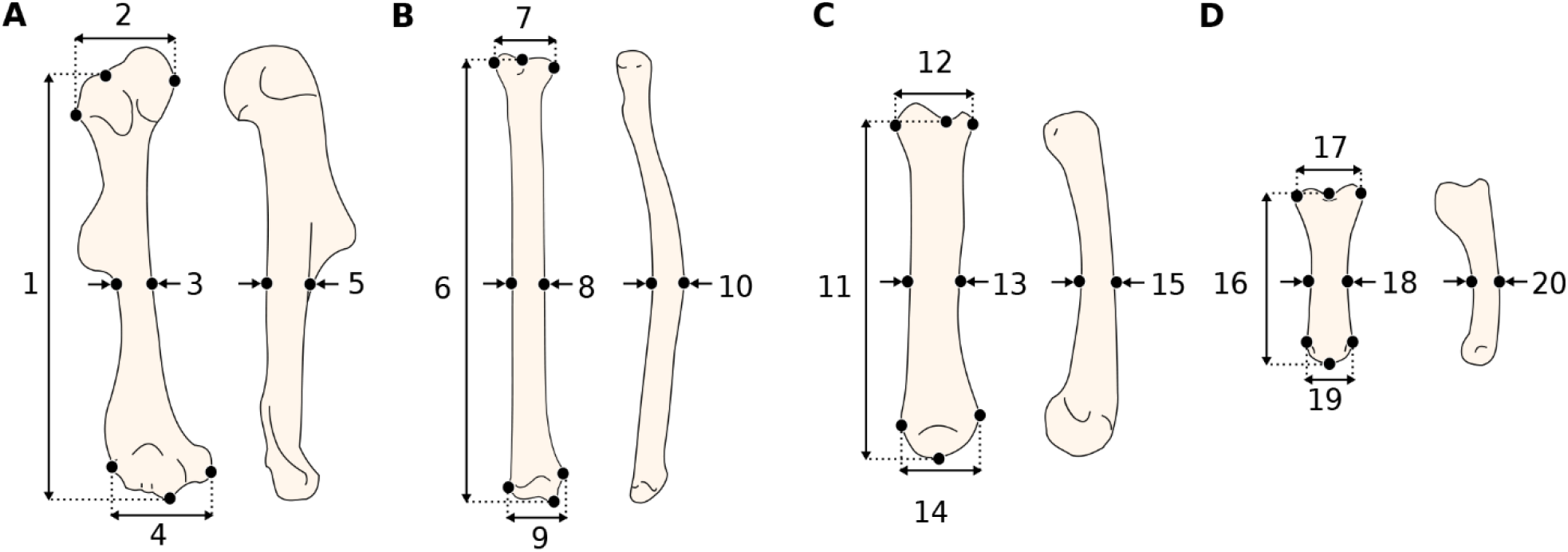
Representation of the linear measurements obtained of the forelimb elements. **A)** Humerus in anterior (right) and lateral (left) view: 1) length, 2) proximal width, 3) mid-shaft width, 4) distal width and 5) height. **B)** Radius in anterior (right) and lateral (left) view: 6) length, 7) proximal width, 8) mid-shaft width, 9) distal width and 10) height. **C)** Third metacarpal in dorsal (right) and lateral (left) view: 11) length, 12) proximal width, 13) mid-shaft width, 14) distal width and 15) height. **D)** First phalanx of the digit III in dorsal (right) and lateral (left) view: 16) length, 17) proximal width, 18) mid-shaft width, 19) distal width and 20) height. Detailed description of each measurement can be found in Table S2.

## Results

The evolutionary model that better predicts the pattern of evolution for all bones measured is the Ornstein-Uhlenbeck (OU) process (Table S3 of SI 2). Therefore, we simulated trait evolution under an OU process on 100 datasets in order to account for error and obtain the results described below.

### Morphological diversity

We inferred morphological diversity for each bone using the determinant and the trace of the simulated trait matrices. Determinants and traces of matrices offer different but complementary generalized metrics to describe the variation of multidimensional data. The matrix trace provides information about the accumulated trait variance, whereas the determinant provides information about the volume occupied by the multivariate data. Both show similar patterns, in which morphological variation increases along the proximo-distal axis, consistent with the timing of limb condensation during development (Figure 3A and B). The early-condensing humerus is the least variable structure, and the late-condensing phalanx is the most diverse element measured, followed by the third metacarpal. All pairwise comparisons between elements are significant (Table 1), although the differences of the determinant distributions of the radius and the metacarpal (*P* = 0.017) are smaller than when using the trace results (*P* < 0.001).

**Table 1.**
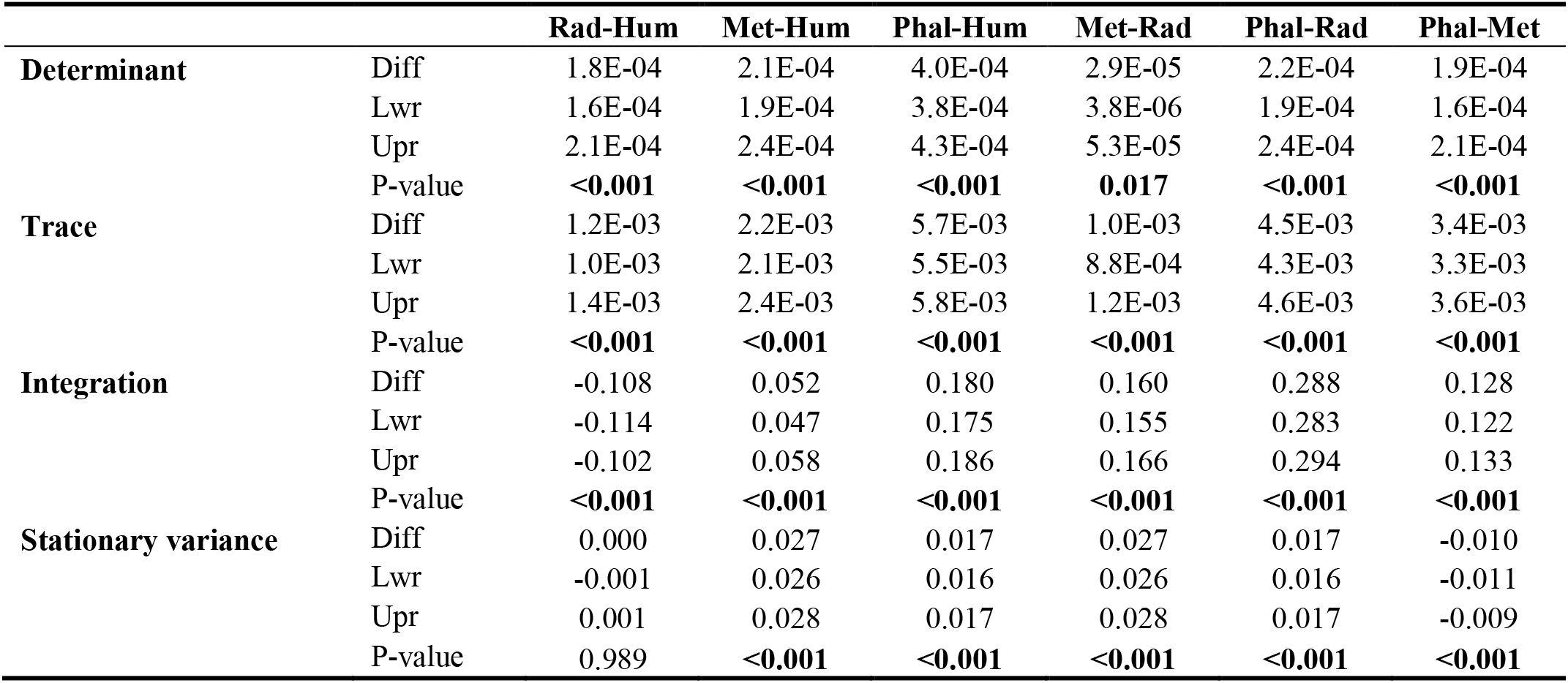
Limb bone pairwise comparison of integration, determinant, trace and stationary variation computed by Tukey Test. Pairwise differences (Diff) of each metric are indicated with the lower (Lwr) and upper (Upr) 95% CI, as well as the adjusted P-values. Hum= Humerus, Rad= Radius, Met= Metacarpus and Phal= Phalanx.

**Figure 3.**
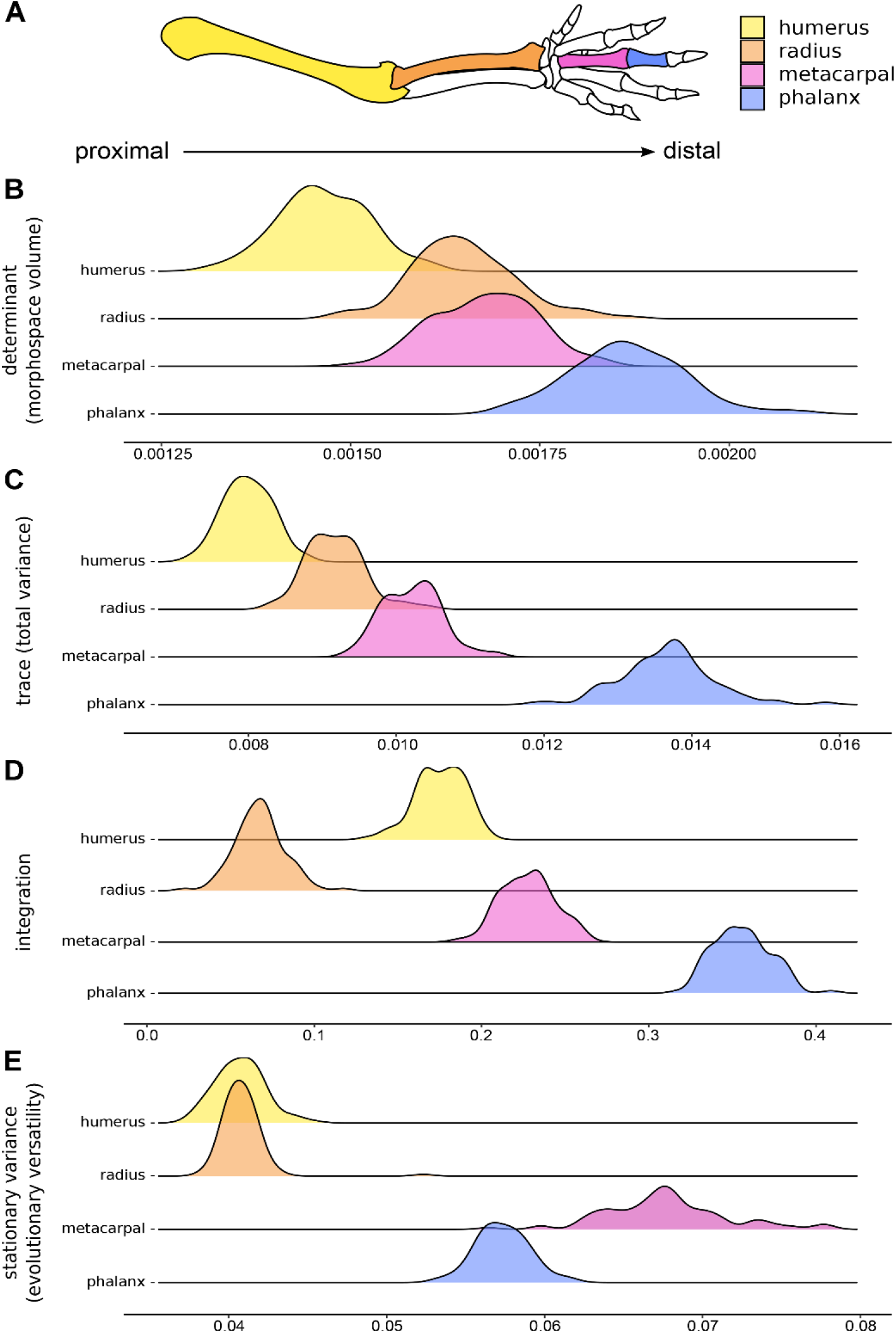
Components of the morphological evolution of forelimb skeletal elements. **A)** Forelimb schematic, with colours indicating bones along the proximo-distal axis: the humerus (yellow), radius (orange), third metacarpal (pink), and the first phalanx of digit III (blue). **B)** Morphological diversity of limb bones inferred by matrix determinant. **C)** Morphological diversity of limb bones, inferred by matrix trace. **D)** Trait integration. **E)** Stationary variance.

### Phenotypic integration

Integration, inferred here by the values of eigenvalue dispersion, is stronger for distal elements compared to proximal ones, the phalanx being the most integrated element, followed by the metacarpal (Figure 3 A and C). The values of integration do not progressively increase along the proximo-distal axis. Instead, the radius is the least integrated structure, and the more proximal humerus is the second least integrated trait. All pairwise comparisons between elements are significant (Table 1).

### Stationary variances

Traits evolving under an OU process exhibit attraction (α) towards their respective adaptive optima (θ). Therefore, we interpreted the tempo of evolution of traits considering the mean stationary variance of each bone, which represents the expected variation when the process is in equilibrium and summarizes the relative influence of stochastic factors in evolution (see Friedman et al., 2021; Gearty et al., 2018; Hansen, 1997; Joly et al., 2018). The stationary variances are significantly higher for distal elements compared to proximal ones. The metacarpal shows the highest stationary variance, followed by the phalanx (Figure 3 A and D). There are no significant differences in the stationary variances at which the humerus and the radius evolve, these values being significantly lower than those of the two autopod elements (Table 2).

## Discussion

The remarkable diversity of limb morphologies seen in mammals reflects the rich ecological and functional diversity that has evolved in this group (Polly, 2007). However, such outstanding morphological diversity does not evolve uniformly among all limb segments. Here, we show a general pattern of limb diversity in Mammalia in which distal elements such as phalanges and metacarpals are in general more disparate and show greater evolutionary versatility than more proximal elements such as the humerus and radius. Our results are consistent with the hypothesis that the timing of element condensation during development regulates the outcomes of morphological evolution. We show that morphological diversity increases along the proximodistal axis with the phalanx being the most variable structure in the mammalian limb, and the humerus being the least diverse element. Conversely, developmental constraints imposed by early versus late development do not seem to determine segment integration; we found that the latest-condensing elements, in the hand, are more integrated than the earlier-condensing humerus and radius. We further show that distal elements evolve, on average, with greater stationary variances than the proximal limb elements. Although we did not find a correspondence of temporal developmental constraints with the integration of structures, the latter do seem to play an important role in morphological diversification.

### Timing of condensation explains the outcome of morphological diversity but not the integration of limb elements

Developmental constraints consist of intrinsic components of the developmental system that may bias the production of phenotypic variability (Wagner, 1988). Consequently, strong constraints have the potential to canalize or restrict the array of phenotypes upon which natural selection can act on, ultimately influencing the course of morphological evolution (Raff, 1996). We here describe a generalized pattern of limb evolution in mammals, in which variation of form increases gradually along the proximo-distal axis. This pattern is consistent with the prediction that lower proximal diversity might be driven by developmental canalization (Hallgrímsson et al., 2002), which, if true, suggests that intrinsic and strong developmental influences on forelimb evolution are shared across Mammalia. Although previous studies have described the outstanding meristic variation in the autopod in contrast with the proximal and intermediate limb (Holder, 1983), here we confirm that such diversity is also detected in the form of hand bones.

The evolutionary variation in development has been proposed as a major determinant of morphological diversity between clades (Cheverud, 1984; Hall, 2012; Wagner, 1988; Watson et al., 2014). Marsupials exhibit less morphological diversity when compared to placentals, for example, they have never evolved fully aquatic life-styles or performing active flight (as whales and bats, respectively). One hypothesised explanation refers to developmental constraints.

Specifically, marsupials are born without being fully developed and need to move from the uterus to the mother’s teat using their forelimbs (Gemmel et al., 2002). An alternative explanation has also been proposed, in which marsupials also had less biogeographical opportunity to diversify into as many ecological niches as placentals (Sánchez-Villagra, 2013). Developmental constraints, ecological limitations, or both in combination therefore are believed to have restricted not only the morphological diversity of the forelimbs of marsupials, but also of their skulls and the jaws (Bennett and Goswami, 2013; Cooper and Steppan, 2010; Fabre et al., 2021; Pevsner et al., 2022; Sánchez-Villagra, 2013; Sears, 2004). Likewise, the within-species variation of forelimb morphology is more integrated in some marsupial species compared to placentals, possibly also reflecting developmental restrictions on these traits in marsupials (Bennett and Goswami, 2011; Kelly and Sears, 2011). Contrary to the expectations of our developmental hypothesis here, however, integration is not greater in early-condensing segments across mammals, but rather is stronger in the later formed metacarpals and phalanges.

The high integration detected in the mammalian hand may reflect the evolutionary modularization of the autopod relative to the proximal limb. This may result from the developmental origin of the autopod, which develops from a conserved module that patterns the relative sizes of phalanges in a range of proportions from a nearly equal-sized pattern to a large-to-small gradient among adult individuals (Kavanagh et al., 2013; Young et al., 2015). Such patterns reflect the dynamic and modular characteristic of limb development, implying that although segments share underlying genetic networks, each bone is mediated by unique markers that attribute their own developmental identity (Cooper et al., 2011; Petit et al., 2017; Schneider and Shubin, 2013; Tanaka, 2016). The modular nature of the limbs likely also explains the occurrence of strong evolutionary integration between anterior and posterior homologous segments, especially in quadrupedal species whose fore- and hind limbs perform similar functions (Botton-Divet et al., 2018; Martín-Serra et al., 2015; Schmidt and Fischer, 2009; Young and Hallgrímsson, 2005). For instance, a reduction in the covariation between homologous regions is observed in the presence of locomotor specialization (Hanot et al., 2017; Martín-Serra et al., 2015; Schmidt and Fischer, 2009; Young and Hallgrímsson, 2005). In bats, for example, the specialization of the forelimb for flight explains the weak degree of integration between anterior and posterior segments, but a high signal of modularity is detectable among elements belonging to the same limb (Young and Hallgrímsson, 2005). The evolutionary divergence of fore- and hindlimb morphologies also involves the divergence of their genetic and developmental factors (Cooper et al., 2012; Farnum et al., 2007), providing relative modularization of hands from feet. On average, we found a strong integration in hand bones of mammals that may reflect such strong modularization of the anterior autopod shared by some taxa. Ultimately, the modularization of the autopod might have facilitated the evolution of functionally specialized traits in mammals. A detailed investigation of how limb integration manifests between clades might elucidate if such patterns are more evident in some groups than others.

The relationship between integration and morphological variation is not always consistent among traits and taxa (Felice et al., 2018). Whereas some studies have shown clear positive associations between high integration and phenotypic variation (Fabre et al., 2021, 2020; Randau and Goswami, 2017), negative associations have been also reported (Felice and Goswami, 2018; Goswami and Polly, 2010). We find no evidence for a strong correspondence of integration with morphological diversity in proximal forelimb segments: the radius exhibits greater diversity of form than the humerus, but presents the weakest values of integration among the bones measured. For the distal elements, however, our results show that the highly integrated autopod, especially the phalanx, also corresponds to the most diverse structure of the limb. These differences might reflect how selection interacts with the intrinsic constraints of variation. Though integration may constrain the evolution of the phenotype to a limited portion of morphospace, it may also promote variation by driving the evolution of these traits in response to selection for functional specialization (Felice et al., 2018; Goswami et al., 2014; Hansen and Houle, 2008; Lande, 1979). Such dynamics appear to be observed in distal elements: high integration in the phalanx and metacarpus, possibly favoured the evolution of functionally specialized autopod structures, contributing to the high variation observed in mammalian hand bones. Future studies will benefit from including extinct taxa, based on fossils, to understand how morphological diversity and integration of limb bones evolved in the deep time, whether these patterns are consistent between major taxonomic and ecological groups and through time, and when they first appeared during mammalian ancestry.

### Evolution of the autopodal elements: functional associations

Functional variation is often a good predictor of the morphological variation of limb bones (Chen and Wilson, 2015; Fabre et al., 2013; Grossnickle et al., 2020; Weaver and Grossnickle, 2020).

Although the distal portion has been suggested to be the most variable structure of the limb, few studies have quantified the functional relationships driving autopod variation in mammals (Almécija et al., 2015; Rolian, 2009; Weisbecker and Schmid, 2007; Weisbecker and Warton, 2006). The hand interacts directly with the surrounding environment, performing important activities such as providing support to the body during locomotion and, in some cases, digging, handling food, grooming and mediating social interactions (Biewener and Patek, 2003; McGrew et al., 2001; Naghizadeh et al., 2020; Weisbecker and Warton, 2006). In our study, we show that the autopod is not only the limb segment showing the highest variation in form, but also that it evolves, on average, with greater stationary variances around their optima than the stylopod and the zeugopod. We suggest that strong functional selection (resulting from the direct impact autopod structures on locomotor performance) combined with the higher potential of development to generate variation in the morphology of more distal limb elements, facilitate the evolution of high autopodial disparity in response to varying environmental demands across mammals.

Examples of autopodial specialisation are widespread among mammals. For example, notable transformations in the metacarpal and phalangeal morphology are observed in cursorial taxa that present specializations allowing for endurance running, typically involving the elongation of the distal limb in relation to proximal segments (Polly, 2007). Morphological adaptations to cursoriality mostly encompass the modification of autopod posture to digitigrady (animals that stand on the distal ends of metapodials and middle phalanges, such as cats and dogs) and unguligrady (animals that stand on their hooved distal-most phalanx, such as horses and cows; Clifford, 2010; Polly, 2007; Wang, 1993). Digitigrady is observed in many carnivorans providing limb elongation and thus increasing stride length (Polly, 2007; Wang, 1993). Extant horses exhibit one of the most dramatic modifications of the third metapodial and phalanges among all unguligrade taxa: the limb is uniquely supported by the third toe, which is considerably enlarged and elongated, whilst the lateral fingers are markedly reduced (McHorse et al., 2019). One recent study suggested that the evolutionary transitions in foot and hand postures are associated with strong selection for rapid changes in increasing body size (Kubo et al., 2019). Although a digital posture presumably implies morphofunctional specialization of the distal limb, it is not clear if the acceleration of body mass evolution during autopod posture transitions has also affected the rates of morphological change of the hand and foot. Autopodial specialisations are also evident among smaller-sized mammals. For example, body size is positively associated with the tempo of evolution of postcranial morphology (hand and foot bones included) in both ground and tree dwelling animals, where medium-sized animals tend to exhibit higher stationary variances than small-sized species(Weaver and Grossnickle, 2020). Nevertheless, in both cases, functional specializations related to the locomotion likely played a role on driving the morphological evolution of the limb, potentially involved with the accelerated evolution of hand bone morphologies. Further investigations are needed to better understand the associations of functional variation with the evolutionary dynamics of limb diversification.

## Conclusion

This study uses a macroevolutionary framework to compare, for the first time, the general patterns of form diversification of proximal and distal limb elements in mammals. Our results reveal that the evolution of the mammalian forelimb involves different patterns of morphological diversification when comparing limb segments along a proximal–distal gradient. We detected that the diversification of autopodial elements was much more dynamic than those of the zeugopod and stylopod, involving higher morphological diversity, stronger integration and greater evolutionary versatility at distal structures. Specifically, we corroborate the premise that the late-condensing distal elements such as metacarpals and phalanges (in the autopod) exhibit higher morphological diversity than early-condensing, more proximal, elements. This pattern might emerge from different levels of constraints during the developmental succession. However, such temporal constraints of development do not explain the patterns of limb evolution alone, as functional specializations may also play an important role on form diversification. Particularly, strong integration at autopod elements might reflect the modularization of hand structures in response to a plethora of functional demands. We highlight the importance of considering known variation during the development to understand the macroevolutionary outcome of adult morphologies and we hope that these results will contribute to better understand the association of limb segment variation with ecological diversity.

## Material and Methods

### Taxonomic sampling and data acquisition

We sampled 638 species of mammals (670 specimens), representing 598 genera of 138 living families (Figure 1). Sampling varies from one to four individuals per genus. We provided micro-CT-scans and surface scans of 58 small to medium sized-specimens from different institutions (available online at MorphoSource.org, Table S1 from SI 1), 23 of them previously used by Martín-Serra and Benson 2020. The digital dataset was combined with 351 meshes available on MorphoSource.org (Table S1 from SI 1). Image stacks were converted into three-dimensional models using Avizo 8.1.1 (1995-2014 Zuse Institute Berlin), where scale dimensions were incorporated based on the voxel size of each scan. Data collection from the digital models was also conducted in Avizo 8.1.1 (1995-2014 Zuse Institute Berlin). We complemented this dataset with measurements provided by caliper of 261 medium to large body-sized species from the mammal collection of the Muséum National d’Histoire Naturelle (Paris, France) (Table S1).

We measured 20 linear distances from anterior limb bones, including the humerus, the radius, the third metacarpal and the first phalanx of digit III. We acquired five measurements for each element: length, widths (proximal, mid-shaft and distal) and height (Figure 2, see detailed description in Table S2 from SI 2). We opted not to include the ulna because this bone is fused to the radius in many taxa (see Sears et al. 2007), preventing the acquisition of such measurements. The metacarpal and first phalanx of digit III were sampled because this is the only digit present in the hands of all mammalian lineages, even in groups that exhibit digit loss or fusion with other autopodial elements, such as in golden moles and ungulates (Clifford, 2010; McHorse et al., 2019; Prothero, 2009). Each individual was measured twice with the subsequent calculation of the mean and standard error in order to verify measurement error. Body mass estimates were assembled from the PanTHERIA database (Jones et al., 2009) and complemented by literature sources when necessary (Table S1 from SI 1). Species taxonomy followed the Mammal Diversity Database published by Burgin et al. (2018).

### Comparative analyses

Analyses were implemented in R 4.1.2 (R Core Team, 2021). We used the phangorn R package (Schliep, 2011) to estimate a maximum clade credibility (MCC) tree from a posterior sample of 10,000 trees published by Upham et al. (2019). Because the incorporation of some species was available only at the genus level, we pruned the MCC tree to genus level, according to the taxa sampled by our study.

Allometry generally explains most part of morphological variation, as body parts usually grow together, masking variation mediated by local development (Marroig, 2007; Raff, 1996). Because we are particularly interested in understanding morphological constraints imposed by the local development of the limb, we decided to remove the allometric component of our dataset in order to reduce variation associated with other sources of development. The genus means of each trait were scaled to size in a linear regression model. First, we transformed body mass into linear scale by taking the cube root prior to log10-transformation (Harmon et al., 2010). We calculated the geometric means of all measurements acquired, including the linear scaled body size, and then we fitted the log10-transformed trait means in a phylogenetic generalized least-squares (PGLS) using the geometric means as a predictor. We grouped the traits by bone and fitted the linear models for each skeletal unit with mvgls() function from mvMORPH R package (Clavel et al., 2019, 2015). We calculated the fit of three models of evolution using LASSO penalization: Brownian Motion (BM), Ornstein-Uhlenbeck (OU) and Early Burst (EB). We compared the likelihood of the model fits with Generalized Information Criterion (GIC).

The OU model of evolution had the best fit for all the linear regressions accounting for body mass using the MCC tree (Table S3 of SI 2). Thus, we simulated trait evolution under OU process on 100 datasets, in order to account for error and conduct the downstream analyses (function mvSIM() from mvMORPH; Clavel et al. 2015, 2019). We repeated the body mass PGLS for the simulated data and calculated the residual covariance phylogenetic matrices.

### Morphological diversity and phenotypic integration

Morphological diversity for each bone was interpreted as the values of the determinant and the trace of simulated matrices. We scaled the determinants by transforming their absolute value to the power of one divided by five, which is the number of dimensions of each matrix (that is, the number of measurements). Differences in the determinant and trace between skeletal elements were evaluated by ANOVA followed by Tukey Tests (function TukeyHSD() from stats R package) of the 95% confidence interval (CI).

We calculated the magnitude of integration for each bone separately, based on eigenvalue dispersion in their respective matrices. We transformed the simulated covariance matrices into correlation matrices and provided integration values as the standard deviation of eigenvalues relative to their theoretical maximum (Haber, 2011; Pavlicev et al., 2009). We calculated the integration as the dispersion of the standard deviation of eigenvalues of our trait matrices, following Pavlicev et al. (2009). For instance, highly integrated traits have most of the independent variance concentrated in the first few eigenvalues, while uncorrelated traits have the variance similarly distributed between eigenvalues (Pavlicev et al., 2009). Eigenvalue dispersion was inferred from CalcEigenVar() function of evolqg R package (Machado et al., 2019; Melo et al., 2015), which calculates the relative eigenvalue variance of the matrix as a ratio between the observed variance and the theoretical maximum for a matrix of the same size and trace (Machado et al. 2019). Differences between distributions were computed by an ANOVA and detailed by Tukey Tests of the 95% CI.

### Macroevolutionary patterns

To assess variability due to the tree topology and branching times, we replicated the body mass linear regressions with 100 trees from Upham et al. (2019). We fitted these linear regressions under OU process, which showed the best support in the previously described PGLS using the MCC tree, and estimated the average rates of evolution (σ^2^) per bone. Because OU is a stochastic process that models the evolution of traits towards an optimum θ with an attraction α, we cannot disentangle the effects of σ^2^ and α to understand the tempo of trait evolution (Hunt, 2012). Therefore, we additionally calculated the mean stationary variance of bones (σ^2^/2α) of OU fitted matrices in order to summarize the relative influence of stochastic factors in evolution (Friedman et al., 2021; Hansen, 1997). We compared their distributions using ANOVA followed by a 95% confidence interval Tukey Test.

## Data availability

Data and codes will be made available on Dryad Digital Repository upon to manuscript publication.

## Acknowledgements

We thank the following collection managers at the MNHN, Paris, for their support during data acquisition: Alexander Nasole, Aude Lalis, Aurélie Verguin, Céline Bens, Géraldine Veron, Jacques Cuisin, Joséphine Lesur and Violaine Colin. Creation of datasets accessed on MorphoSource was made possible by the following funders and grant numbers: NSF DBI-1701713, 1701714, 1701737, 1702263, 1701665, 1701767, 1701769, 1701870, 1701797, 701851, 1702442, 1902105, BCS 1317525, BCS 1540421, BCS 1552848; ERC-2015-STG-677774 (TEMPO Mammals project to RB) and Leakey Foundation. Project funding for digital acquisition of each used specimen is detailed in Table S1 of Supporting Information 1. We also thank Anjali Goswami, Helder Gomes Rodrigues, Eric Guilbert and Loïc Kéver for their insightful comments during the elaboration of this study. This work was supported by CNPq doctoral grant to PSR (process #204841/2018-6).

## Author contributions

Conceptualization: PSR, ACF and AH; data curation: PSR and RB; data acquisition: PSR; methodology: PSR, ACF, JC and AH; analyses design: PSR and JC; result interpretation: PSR, ACF, JC, AH; writing original draft: PSR; review and editing: PSR, ACF, RB, AH. All authors approved the manuscript submission.

## Competing interests

The authors declare no competing interests.

## Notes

### Competing Interest Statement

The authors have declared no competing interest.

